## Supplementary Data 01 for "Cell membrane glycan contents are biochemical factors that constitute a kinetic barrier to viral particle uptake in a protein-nonspecific manner"

| Name | Source | Identifer |
|---|---|---|
| **Lectins** | | |
| Sambucus sieboldiana | J-chemical | J118 |
| ConA | J-chemical | J103 |
| Aleuria aurantia Lectin (AAL),Aleuria aurantia | J-chemical | J101-R |
| Peanut Lectin, Lyophilized | Wako Fujifilm | 165-15031 |
| Wheat Germ Lectin | Wako Fujifilm | 126-02811 |
| DSA Lectin from Datura stramonium(jimson weed, thorn apple) | SIGMA | L2766 |
| Maackia Amurensis Lectin, Lyophilized | wako fujifilm | 139-10891 |
| **Antibodies** | | |
| Anti-MUC1 antibody [EP1024Y] | Abcam | ab45167 |
| Purified anti-mouse/human CD44 | BioLegend | 103001 |
| Purified anti-human CD24 | BioLegend | 311101 |
| Purified anti-human CD164 | BioLegend | 324802 |
| Purified anti-CD49C | BioLegend | 865201 |
| Purified anti-human/mouse CD49f Antibody | BioLegend | 313602 |
| Purified anti-human CD138 (Syndecan-1) | BioLegend | 356501 |
| Monoclonal Antibody to Receptor Tyrosine Protein Kinase erbB-2 (ErbB2), C12 | Cloud-Clone Corp | MAB867Hu22 |
| PLXNB2 Rabbit pAb | ABClonal | A10069 |
| Anti-Angiotensin Converting Enzyme 2, Ectodomain, Human, Goat-Poly <Anti-ACE-2> | R&D systems | AF933 |
| Anti-SARS-CoV/SARS-CoV-2 (COVID-19) Spike, Mouse-Monoclonal antibody (1A9) | GTX | 632604 |
| Goat anti-Rabbit IgG (H+L) Cross-Adsorbed Secondary Antibody, Alexa Fluor™ 555 | Thermo Fisher | A-21428 |
| Goat anti-Rat IgG (H+L) Cross-Adsorbed Secondary Antibody, Alexa Fluor™ 555 | Thermo Fisher | A-21434 |
| Goat anti-Mouse IgG (H+L) Cross-Adsorbed Secondary Antibody, Alexa Fluor™ 555 | Thermo Fisher | A-21422 |
| Anti Mouse IgG CF568 Goat 50uL | BTI | 20100-1 |
| Alexa 647 labeled anti-Mouse IgG secondary antibody | Thermo Fisher | A-21235 |
| **Other chemicals and reagents** | | |
| SNAP –Cell TMR star | New England Biolabs | S9105S |
| SNAP-Surface 488 | New England Biolabs | S9124S |
| SNAP-Surface Alexa 647 | New England Biolabs | S9136S |
| Alexa Fluor™ 488 NHS Ester (Succinimidyl Ester) | Thermo Fisher | A20000 |
| Alexa Fluor™ 647 NHS Ester (Succinimidyl Ester) | Thermo Fisher | A37566 |
| DiIC18(5) solid (1,1'-Dioctadecyl-3,3,3',3'-Tetramethylindodicarbocyanine, 4-Chlorobenzenesulfonate Salt, or DID | Thermo Fisher | D7757 |
| BODIPY™ FL DHPE | Thermo Fisher | D3800 |
| 1,2-Dioleoyl-sn-glycero-3-phosphatidylcholine | Avanti Polar Lipids | 850375 |
| 1,2-dioleoyl-sn-glycero-3-phosphatidylserine | Avanti Polar Lipids | 840035 |
| 1,2-dioleoyl-sn-glycero-3- [(N-(5-amino-1-carboxypentyl)iminodiacetic acid)succinyl] (DGS-NTA(Ni) | Avanti Polar Lipids | 790404 |
| tetramethylrhodamine thiocarbamoyl-1,2-dihexadecanoyl-sn-glycero-3-phosphoeth- anolamine, triethylammonium salt | Thermo Fisher | T1391 |
| Perylene | Sigma | 394475-1G |
| Hoechst 33342 solution | Dojindo | H342 |
| Plasmem Bright Red | Dojindo | P505 |
| MemGlow 590 | Cytoskeleton inc. | MG03-02 |
| Cysteamine (MEA) | Sigma | #30070-50G |
| Glucose Oxidase type seven from Aspergillius | Sigma | #G2133-250KU |
| Catalase, from Bovine Liver | Wako Fujifilm | #035-12903 |
| Lenti-X™ Concentrator. | Takara Bio | 631231 |
| Lenti-X™ p24 Rapid Titer Kit | Takara Bio | 631476 |
| 10x Buffer BXT | iba | # 2-1042-025 |
| Strep-Tactin XT 4Flow high capacity | iba | # 2-5030-002 |
| GST-bind resin | Merck Millipore | # 70541-3 |
| PreScission Protease | Cytiva | # 27-0843-01 |
| Micro Bio-Spin™ P-6 Gel Columns, Tris Buffer | Biorad | 7326221 |
| 6.5 mm Transwell® with 0.4 µm Pore Polyester Membrane Insert, Sterile | Corning | 3470 |
| Uniform Silica Microspheres Non-functionalized Silica | Bangs lab | SS05N |
| **Cell line** | | |
| Calu-3 | ATCC | HTB-55 |
| HEK 293T | ATCC | CRL-11268 |
| **Plasmids** | | |
| pLV-eGFP | Addgene | #36083 |

| | | |
|---|---|---|
| AAV GFP | Addgene | #49055 |
| pAdDeltaF6 | Addgene | #112867 |
| pAAV2/2 | Addgene | #104963 |
| AxCAEGFP | Riken BRC | RDB03343 |
| pCAG-HIVgp | Riken BRC | RDB04394 |
| pCMV-VSV-G-RSV-Rev | Riken BRC | RDB04393 |
| pCMV-SARS-CoV-2 Spike-RSV-Rev | Kaizuka et al. (2023) | NA |
| ACE2-iRFP | Kaizuka et al. (2023) | NA |
| TMPRSS2-TagRFP657 | Kaizuka et al. (2023) | NA |
| SNAP-MUC1(42TR) | This study | NA |
| SNAP-MUC1(0TR) | This study | NA |
| SNAP-MUC1(4TR) | This study | NA |
| SNAP-MUC1(14TR) | This study | NA |
| SNAP-CD43 | This study | NA |
| SNAP-CD44 | This study | NA |
| SNAP-CD24 | This study | NA |
| SNAP-CD164 | This study | NA |
| SNAP-SDC1 | This study | NA |
| SNAP-F174B | This study | NA |
| SNAP-VCAM1 | This study | NA |
| SNAP-VAMP2 | This study | NA |
| SNAP-PD-1 | This study | NA |
| SNAP-EPHB1 | This study | NA |
| CD43-mTagBFP2 | This study | NA |
| PODXL-mTagBFP2 | This study | NA |
| TMEM123-mTagBFP2 | This study | NA |
| EFNB2-mTagBFP2 | This study | NA |
| EPHB1-mTagBFP2 | This study | NA |
| Jag1-mTagBFP2 | This study | NA |
| Dll4-mTagBFP2 | This study | NA |
| GYPC-mTagBFP2 | This study | NA |
| LCK10-mCherry | This study | NA |
| dStrep-SNAP-MUC1(14TR)_ecto-His | This study | NA |
| dStrep-SNAP-CD43_ecto-His | This study | NA |
| pGEX6p1-MUC1(14TR)_ecto-His | This study | NA |
| pGEX6p1-CD43_ecto-His | This study | NA |
| **Software** | | |
| Python | | 3.9.2 |
| Matplotlib | | 3.7.2 |
| Numpy | | 1.22.1 |
| Pandas | | 1.4.2 |
| numpyro | | 0.11.0 |
| jax | | 0.4.8 |
| Fiji ImageJ | | 2.1.0/1.54d |
| Affinity Designer | Serif | 1.10.8 |
| Data Graph | VisualDataTools | 5.2 |
