## Supplementary Data 03 for "Cell membrane glycan contents are biochemical factors that constitute a kinetic barrier to viral particle uptake in a protein-nonspecific manner"

| UniProt Entry | Protein # | Gene name | Protein names | Total Etcodomain (AA) | # of membrane-spanning | Max Etcodomain (AA) | GlycoEP(N,O) | Predicted Glycosylation /AA | Net N/O Glyc | Predicted Glycosylation /AA | Mean (Glyco EP & Net N/O Glyc) |
|---|---|---|---|---|---|---|---|---|---|---|---|
| P15941 | | MUC1 | MUC1 0TR(del84 | 294 | 1 | 294 | | 0.275510204 | | 0.227891156 | 0.25170068 |
| P15941 | | MUC1 | MUC1 4TR(del76 | 374 | 1 | 374 | | 0.267379679 | | 0.229946524 | 0.248663102 |
| P15941 | | MUC1 | MUC1 14TR(del5 | 574 | 1 | 574 | | 0.261324042 | | 0.236933798 | 0.24912892 |
| Q9JI71 | | Dll4 | Delta-like protein | 505 | 1 | 505 | | 0.061386139 | | 0.055445545 | 0.058415842 |
| Q9QXX0 | | Jag1 | Protein jagged-1 | 1033 | 1 | 1033 | | 0.056147144 | | 0.010648596 | 0.03339787 |
