## Supplementary information for "Cell membrane glycan contents are biochemical factors that constitute a kinetic barrier to viral particle uptake in a protein-nonspecific manner"

### Supplementary Figure 1.

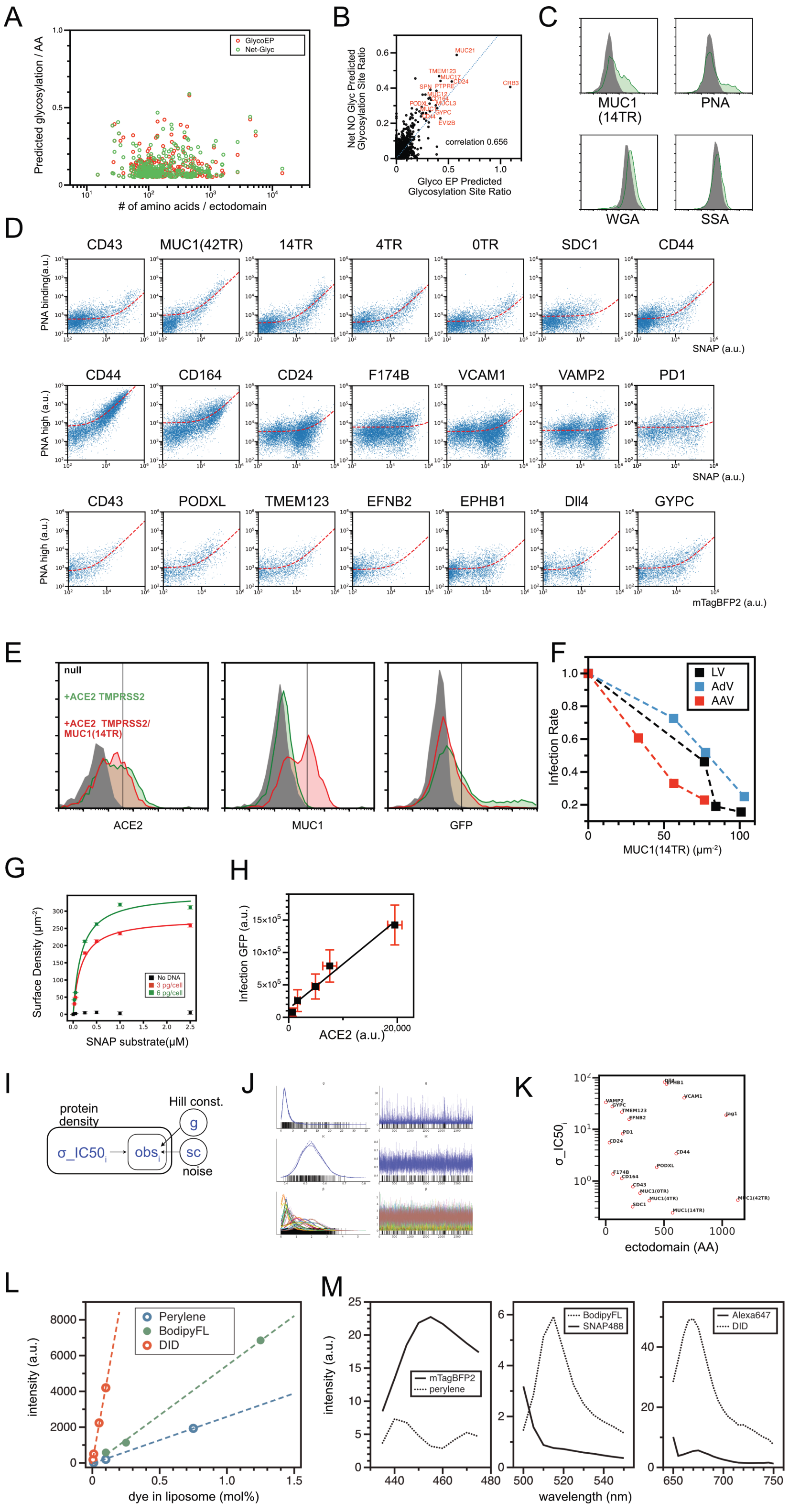

A. Glycosylation site prediction by Net O Glyc/Net N Glyc and GlycoEP. B. Comparisons of the number of predicted glycosylation sites by both software packages. C. Flow cytograms demonstrating the binding of three different fluorescent labeled lectins (PNA, WGA, SSA) to HEK 293T cells expressing MUC1(14TR) proteins. Protein expression probed by SNAP tag fluorescence (upper left), PNA binding (upper right), WGA binding (bottom left), and SSA binding (bottom right) were shown. D. Two-dimensional flow cytograms of PNA binding assay to HEK 293T cells expressing various membrane proteins. Protein density was shown in either SNAP tag (SNAP surface 488) or mTagBFP2 fluorescence. Two different PNA concentrations were used to have wide enough dynamic range in PNA signals to resolve glycosylation in both low- and high- glycosylated proteins. PNA signals between low- and high- PNA concentrations and between SNAP tag and mTagBFP2 were adjusted with data of sample measured in both conditions (CD43, CD44). E. Examples of flow cytograms in virus infection inhibition assays in HEK293T cells and SARS-CoV2-PP. Expressions of ACE2-mTagBFP2 (left) and SNAP-MUC1(14TR) (middle) were measured at the time of infection, and expression of GFP (right) was measured 2 days after the infection in the same batch of cells. Dark is null HEK293T cells without ACE2 and MUC1 expression. Green lines for HEK293T-ACE2 cells without MUC1 expression. Red lines for HEK293T-ACE2-MUC1 cells. F. Infection inhibition assays for HEK293T cells and various other viruses (lentivirus, adenovirus, and adeno-associated virus). Mean GFP expression of infected cells are plotted along with mean MUC1 (14TR) densities in cells at the time of infection. These were measured by flow cytometry, in similar manner as in the assay for SARS-CoV2-PP. G. Measurement of surface densities of SNAP – Surface 488 labeled SNAP – MUC1 (14TR) proteins in HEK 293T cells, transfected with different amount of plasmid DNAs. H. Integrated amount of GFP expression in SARS-CoV2-PP infected HEK 293T cells compared with integrated amount of ACE2-mTagBFP2 expression at the time of infection. I. Graphical model for Bayesian hierarchical sigmoidal regression of infection inhibition assay data. J. Results of 4 repeated rounds of MCMC samplings by Numpyro for inferences for each parameter. K. Relations between amino acid sequence for ectodomain of each protein and molecular specific IC50 density in sigmoidal inhibitory function inferred from Bayesian hierarchical modeling (σ_IC50). L. Flow cytometry measurements for liposomes containing serially diluted dye-conjugated lipids and fluorescent membrane incorporating molecules (Bodipy-FL, peryelene, and DID) with indicated mol%. Linear fitting shown was used for calibration. M. Fluorescence emission spectrum for equimolar molecules (50μM for green and far-red channels, and 100μM for blue channel), excited at 405 nm, 488 nm, and 638 nm, respectively. Membrane dyes were measured as incorporated in liposomes. Purified recombinant mTagBFP2 was used.

### Supplementary Figure 2.

A

| Gene | Predicted Glycosylation Rate | Ectodomain(AA) | Glycans: Molecule/Total(%) |
|---|---|---|---|
| CD44 | 0.199045 | 628 | 11.8046 |
| TMEM123 | 0.438849 | 139 | 2.5065 |
| CD164 | 0.315217 | 138 | 1.6772 |
| TGOLN2 | 0.169916 | 359 | 1.3516 |
| CD24 | 0.484375 | 32 | 1.0585 |
| SDC1 | 0.192641 | 231 | 0.8097 |
| C11orf24 | 0.324538 | 379 | 0.4663 |
| MANSC1 | 0.177374 | 358 | 0.319 |
| SDC3 | 0.257772 | 386 | 0.2607 |
| FAM168B | 0.181818 | 99 | 0.2366 |

B

| Gene | Predicted Glycosylation Rate | Ectodomain(AA) | Glycans: Molecule/Total(%) |
|---|---|---|---|
| ERBB2 | 0.028617 | 629 | 6.505 |
| ITGB1 | 0.034653 | 707 | 2.5156 |
| FAT1 | 0.021039 | 4159 | 2.2665 |
| APP | 0.049048 | 683 | 2.2161 |
| ITGA3 | 0.024008 | 958 | 1.9573 |
| PTPRF | 0.03528 | 1233 | 1.8469 |
| TNFRSF21 | 0.143322 | 307 | 1.4557 |
| MEGF8 | 0.051546 | 2619 | 1.3687 |
| JAG1 | 0.051307 | 1033 | 1.3565 |
| TFRC | 0.026826 | 671 | 1.2298 |
| ITGA6 | 0.022904 | 1026 | 1.2022 |
| ADAM9 | 0.021707 | 668 | 1.19 |
| CELSR1 | 0.054365 | 2447 | 1.1267 |
| CPD | 0.071823 | 1267 | 1.0421 |
| PLXNB1 | 0.059184 | 1470 | 0.9565 |
| CDH1 | 0.042419 | 554 | 0.9493 |
| THSD7A | 0.063182 | 1559 | 0.8305 |
| MET | 0.02591 | 907 | 0.7945 |
| PLXNB2 | 0.026763 | 1177 | 0.7665 |
| DCBLD2 | 0.056399 | 461 | 0.7511 |

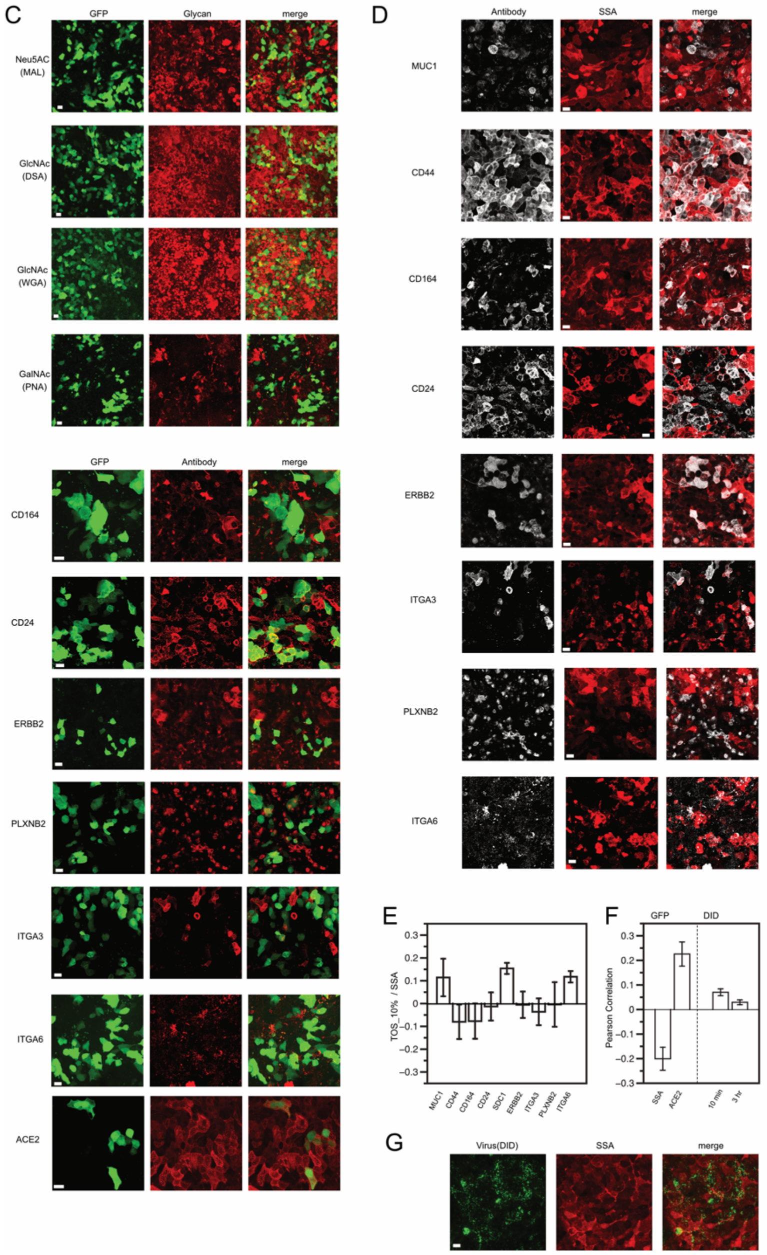

A-B. List of highly expressed membrane proteins in air liquid interface cultured Calu-3 cells, aligned in the order of normalized expression shown in bulk RNA-seq analysis by Wyler E et al. (GSE148729). Ratios for total glycans of each molecule are calculated by combining with predicted glycosylation rate determined in our study and the ratio of expression from the RNA seq result. The panel A is the list for proteins predicted to have predicted glycosylation rate > 0.15, and the panel B is the list for other membrane proteins. C. Maximal projection of z-stack of images. SARS-CoV2-pp infected air liquid interface (ALI) cultured Calu-3 cell monolayer, imaged by the binding of Alexa Flour 647-labeled lectins specific for indicated glycans or the binding of primary and Alexa Flour 555 labeled secondary antibodies against indicated proteins, and the expression of GFP derived from infected viruses. D. Maximal projection of z-stack of images. SARS-CoV2-pp infected air liquid interface (ALI) cultured Calu-3 cell monolayer, imaged by the binding of Alexa647-labeled Neu5AC specific lectin from Sambucus sieboldiana (SSA) and the binding of primary and Alexa Flour 555 labeled secondary antibodies against indicated proteins. E. Correlation in top 10% in both axes in TOS analysis, for colocalizations in SSA and glycoproteins. Error bars are standard error of mean from images from three or more different samples. F-G. DID-labeled SARS-CoV2-pp was incubated on ALI Calu-3 cells for 10 min or 3 h, and the Pearson correlation between DID and SSA was calculated at the pixel level. Correlation values were compared to those of positively and negatively correlated samples (ACE2, SSA, versus GFP, respectively). Example virus and lectin images after 10 min virus incubation are shown in G. Scale bar 20 μm.

**Supplementary Figure 3.**

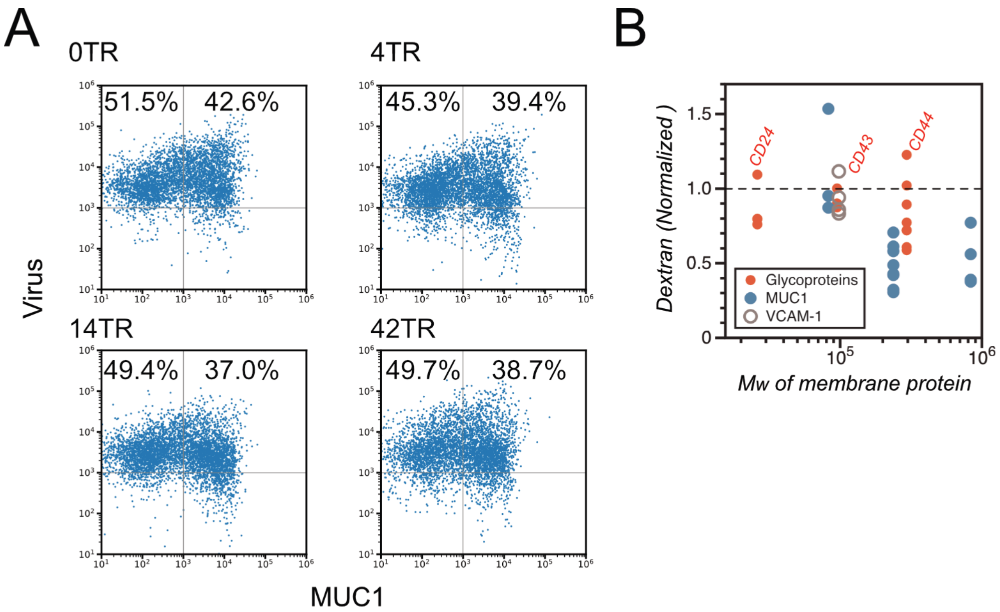


A. Dual-color flow cytogram (DID and SNAP surface 488) of DID-labeled SARS-CoV2-pp-bound HEK293T cells expressing MUC1 with tandem repeat domains of the indicated lengths. MUC1 was labeled and quantified with SNAP surface 488. B. Normalized mean fluorescence intensity of FITC-labeled dextran endocytosed into HEK293T cells expressing the indicated membrane proteins, measured by flow cytometry. Data are normalized to a control sample without exogenous membrane protein expression and plotted against the molecular weight of the introduced membrane protein. Three different MUC1 molecules with different molecular weights (0TR, 14TR, and 42TR) were used. Each dot represents an independent measurement.

**Supplementary Figure 4.**

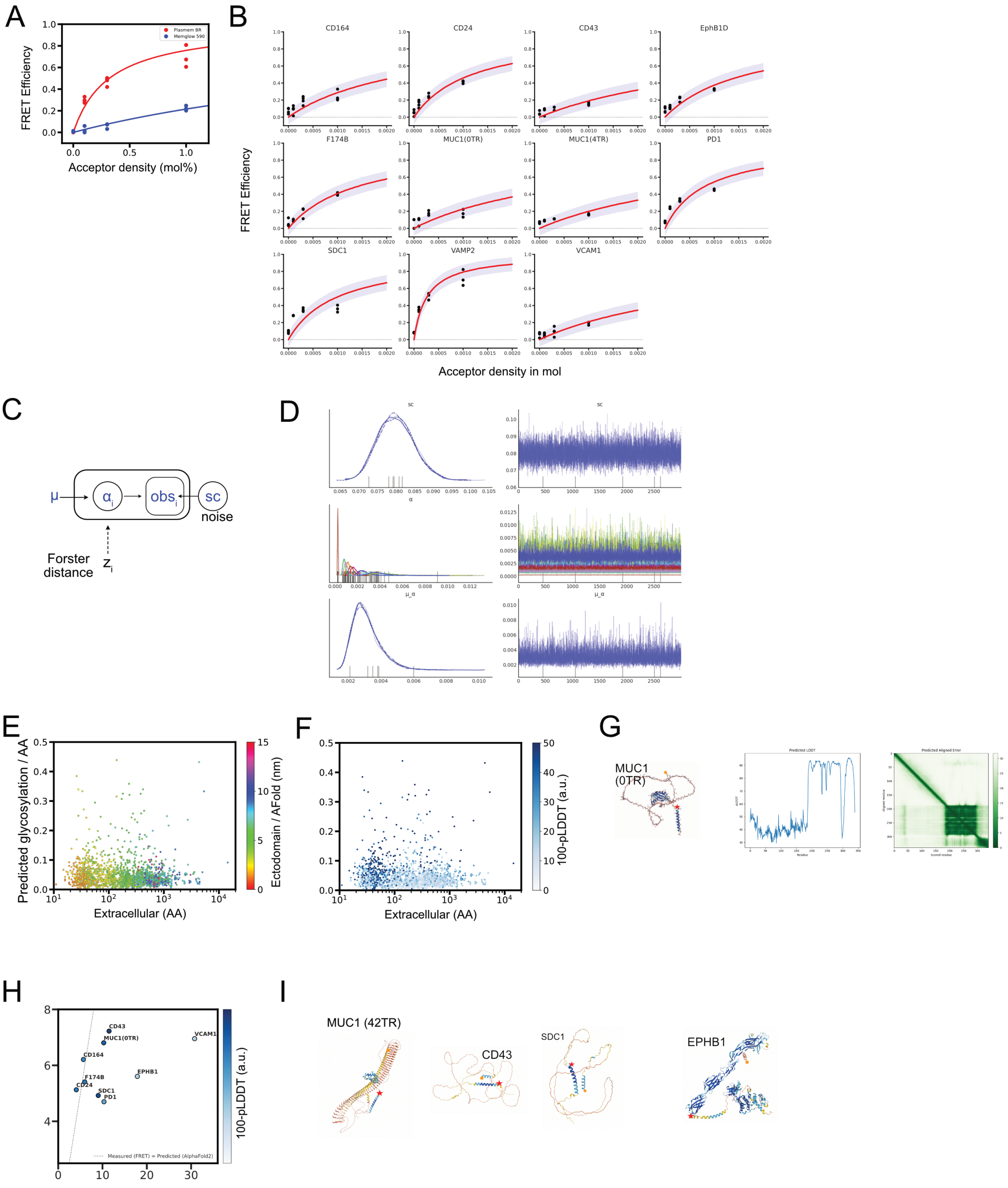


A. FRET efficiency estimated from FLIM imaging for HEK 293T cells expressing SNAP-VAMP2 conjugated with SNAP – Surface 488, where cell membranes contained different mean acceptor densities of either PlasMem Bright Red or MemGlow 590. Lines are learned predicted mean FRET efficiencies from Bayesian hierarchical inference (see method), and purple area represents one sigma below and above the lines. B. FRET efficiency estimated from FLIM imaging for cells expressing each protein at different mean acceptor densities. Lines are learned predicted mean FRET efficiencies from Bayesian hierarchical inference (see method), and purple area represents one sigma below and above the lines. C. Graphical model for Bayesian hierarchical regression of FLIM – FRET data. D. Results of 4 repeated rounds of MCMC samplings by Numpyro for inferences for each parameter. E. Protein size derived from the predicted ectodomain conformations by Alpha fold 2, plotted on Figure 1C by color map. F. Predicted local Distance Difference test scores (pLDDT) for the predicted ectodomain conformations by Alpha fold 2, were subtracted from 100, and were plotted on Figure 1C by color map. By applying this color map that is inverse to original pLDDT score, proteins predicted not to fit well to reference models were highlighted by darker blue mark. G. Alpha Fold 2 (ver 2.2 in Google colaboratory) predicted conformation of MUC1 (0TR), with predicted LDDT along with amino acid and the map of predicted aligned error. H. Relations between pLDDT subtracted from 100 in Alpha Fold 2 prediction and measured Flory radius from FLIM – FRET assays. Dot line indicates where the measured RF is equal to the Alpha Fold 2 predicted length. I. Conformation drawings of proteins predicted by Alpha Fold 2. Yellow and red stars indicate the two ends of ectodomain.

**Supplementary Figure 5.**

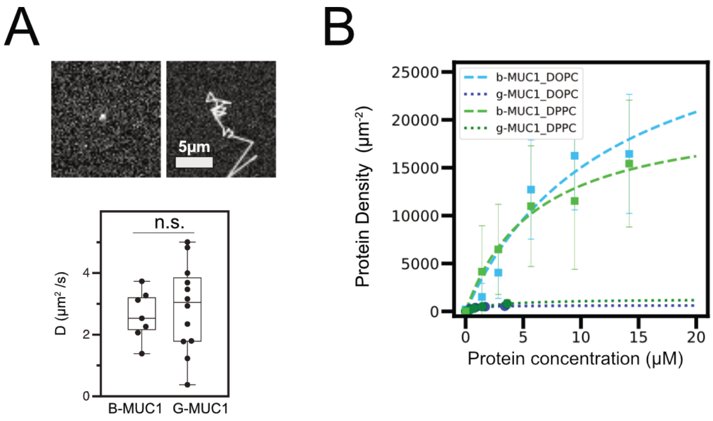


A. An example of a snapshot image of single molecular SNAP–cell TMR star conjugated g-MUC1 (14TR) (upper left) and a tracked path of its diffusive motion on a supported lipid bilayer (upper right). Statistics of diffusion coefficients of g-MUC1 and b-MUC1 calculated from tracked paths (lower). n.s.: nonspecific in a two-sided unpaired t-test with Welch's correction. B. Representative result for flow cytometry analyses of protein binding to lipid bilayer coated silica beads. Bar is standard deviation in each measurement. Lines are regression curves to receptor binding model $Bx/(K_D + x)$, where $x$ is protein concentration, $K_D$ is the dissociation constant and $B$ is the saturated density.

**Supplementary Figure 6.**

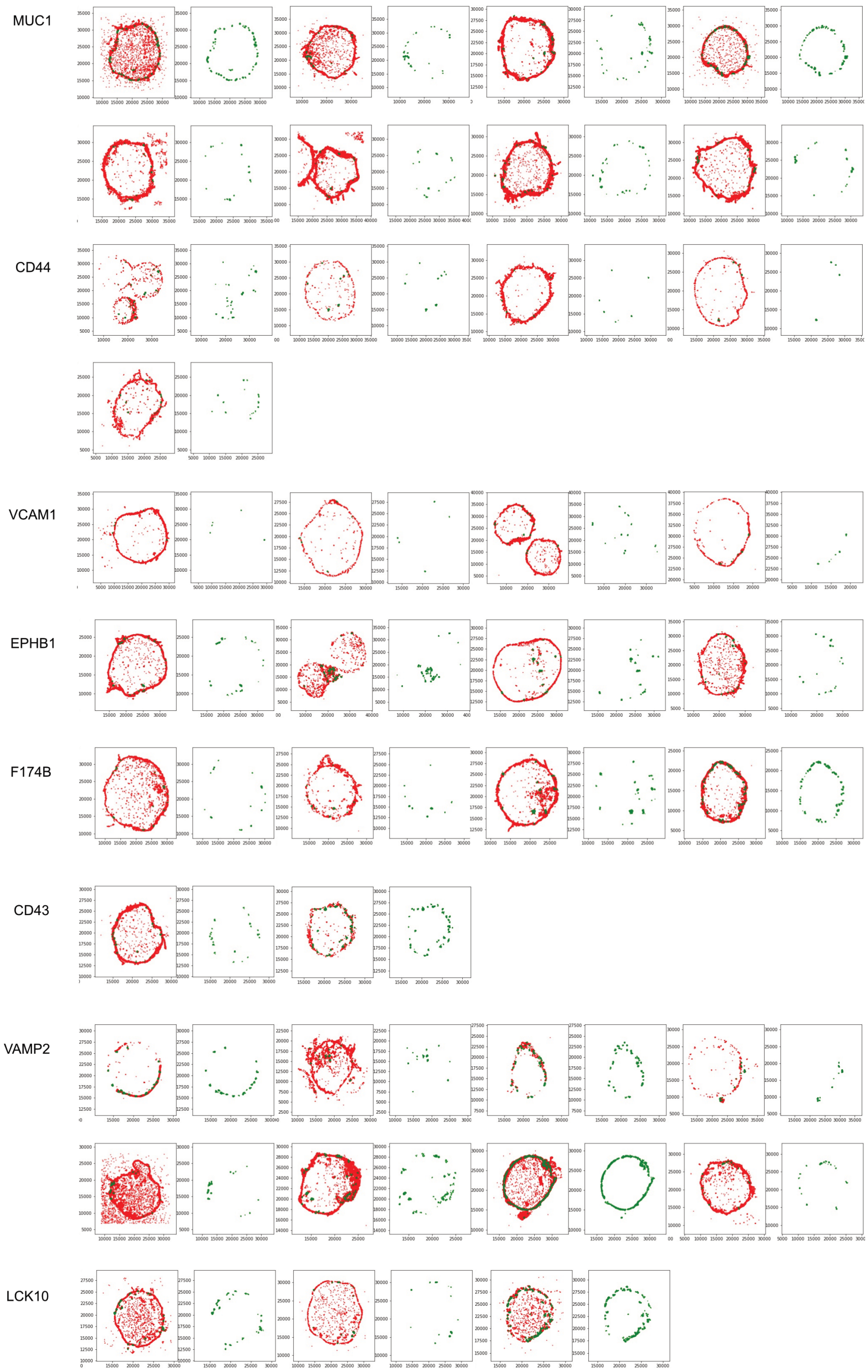


STORM images of all analyzed cells, expressing designated proteins. The detected spots of SNAP-surface Alexa 647 bound to each membrane protein are shown in red, and the spots of CF568-conjugated anti-mouse IgG secondary antibody that recognizes Spike on SARS-CoV2-PP are shown in green. For cells, a pair of two-color composite images and a CF658-only image are shown. mNumbers on axes are coordinates in nanometer.

**Supplementary Data 1.**

List of all reagents and software used in this study.

**Supplementary Data 2.**

List of 2515 human membrane proteins whose ectodomain amino sequences were used for glycosylastion predictions by NetN/O Glyc and GlycoEP. Length of amino acid sequences in ectodomains as well as ratio of glycosylation site per amino acid are listed for all molecules.

**Supplementary Data 3.**

Glycosylation prediction data for MUC1 truncation mutants and mouse proteins used in this study.
